# Supplementation in vitamin B3 counteracts the negative effects of tryptophan deficiencies in bumble bees

**DOI:** 10.1101/2022.06.21.497066

**Authors:** M.L Tissier, S. Kraus, T. Gómez-Moracho, M. Lihoreau

## Abstract

Increasing evidence highlights the importance of diet content in 9 essential amino acids for bee physiological and behavioral performance. However, the tenth essential amino acid, tryptophan, has been overlooked as its experimental measurement requires a specific hydrolysis. Tryptophan is the precursor of serotonin and vitamin B3, which together modulate cognitive and metabolic functions in most animals. Here, we investigated how tryptophan deficiencies influence the longevity and behavior of bumble bees. Tryptophan-deficient diets led to a moderate increase in food intake, aggressiveness and mortality compared to the control diet. Vitamin B3 supplementation in tryptophan-deficient diets tended to buffer these effects on food intake and survival, and significantly reduced workers aggressiveness. Considering that the pollens of major crops and common plants, such as corn and dandelion, are deficient in tryptophan, these effects could have a strong impact on bee populations and their pollination service.

## Introduction

Tryptophan is one of the 10 essential amino acids (EAAs) in all eukaryotes and even in some prokaryotes (Baker, 2008; de Groot, 1953; Kantak et al., 1980; Meisinger and Speer, 1979; Walz et al., 2013). It is the precursor of serotonin, melatonin, and vitamin B3 (niacin) (de Arruda et al., 2013; Kohlmeier, 2015). As a precursor of serotonin, tryptophan modulates aggressive behavior in most vertebrates and invertebrates studied so far (Bubak et al., 2020). As a precursor of vitamin B3, tryptophan is also involved in cell respiration and ATP synthesis. Indeed, vitamin B3 can be decomposed into two molecules, nicotinic acid and nicotinamide, used for the synthesis of the coenzyme nicotinamide adenine dinucleotide (NAD) in vivo (Kohlmeier, 2015; Wan et al., 2011) which is indispensable for the effective functioning of the Krebs cycle in the mitochondria.

Animals cannot synthesize tryptophan, and for most species B3 synthesis from tryptophan is poorly efficient (Baker, 2008). Therefore, both tryptophan and vitamin B3 must thus be obtained daily from food. Diet deficiencies in tryptophan and its derivates can lead to dementia, diarrhea and dermatitis (i.e. skin rashes) in humans (Hegyi et al., 2004; Wan et al., 2011), the black-tong syndrome in dogs (Baker, 2008), and aggressiveness and growth retards in rats (Kantak et al., 1980; Krehl et al., 1945; Walz et al., 2013). Similar effects are also observed in wild animals. For instance, a lack of nicotinamide causes high rates of maternal infanticides in the farmland European hamster restricted to monotonous diets of corn, leading to a major reduction in its reproductive success (Tissier et al., 2017). Although there is an abundant literature in vertebrates (Baker, 2008; Hegyi et al., 2004; Kantak et al., 1980; Krehl et al., 1945; Meisinger and Speer, 1979; Tissier et al., 2017; Walz et al., 2013), studies of these effects on invertebrates are scarce (Bubak et al., 2020; de Groot, 1953).

Malnutrition is a leading cause of bee decline (Klein et al., 2017; Vanbergen and Initiative, 2013). The generalization of crop monocultures constrains pollinators to suboptimal monotonous diets (Goulson et al., 2015; Klein et al., 2017). For bees that rely on carbohydrates in nectar, and proteins, minerals, vitamins, and lipids in pollen (Brodschneider and Crailsheim, 2010; Vaudo et al., 2016), inappropriate nutrient intake can strongly impair their physiology (Conroy et al., 2016; Di Pasquale et al., 2013), survival (Conroy et al., 2016), and colony growth (Moerman et al., 2017). Recent studies show the concentration and balance of key nutrients in nectars and pollens (e.g. content in EAAs (Leonhardt and Blüthgen, 2012; Moerman et al., 2017; Moerman et al., 2016; Stabler et al., 2015; Vanderplanck et al., 2014) or omega 6:3 ratio (Arien et al., 2015)) can influence many parameters related to bee performances, such as olfactory and tactile associative learning (Arien et al., 2015), egg production (Vanderplanck et al., 2014), and larval development (Moerman et al., 2017; Moerman et al., 2016; Vanderplanck et al., 2014). Regarding protein quality, studies are increasingly looking at the total concentration in total amino acids or EAAs (Leonhardt and Blüthgen, 2012; Moerman et al., 2017; Moerman et al., 2016; Ryder et al., 2021; Stabler et al., 2015; Vanderplanck et al., 2014) as an index of diet quality to which colony growth has been associated (Leonhardt and Blüthgen, 2012; Moerman et al., 2017; Moerman et al., 2016).

Tryptophan has been overlooked in those studies (Kriesell et al., 2017; Leonhardt and Blüthgen, 2012; Moerman et al., 2017; Moerman et al., 2016; Ryder et al., 2021; Vanderplanck et al., 2014) as its experimental measurement requires a specific alkaline hydrolysis in contrast to all others essential amino acids necessitate an acid hydrolysis (Standifer et al., 1980). Early work indicates tryptophan deficiencies reduce growth and survival in the Western honeybee (de Groot, 1953). A diet content of 12 mg of tryptophan per gram of food is considered optimal in honey bees, maximizing workers food intake, antioxidant capacity, serotonin concentration, hypopharyngeal gland acinus size (the primary organ secreting jelly), body weight and lifespan. Below and above this concentration detrimental effects were recorded, such as the reduction of food intake and lifespan (Fengkui et al., 2015). While this is an important first step, virtually nothing is known on the effects of deficiencies in tryptophan or its derivates on other major pollinators in the context of widespread decline. While vitamin B3 is one of the most abundant B-complex vitamins found in pollen collected by bees (de Arruda et al., 2013), pollen, leaves and seeds of many common plants are known to be deficient in tryptophan and to contain a bounded form of vitamin B3, non-bioavailable to animals. These include most cereals and some weeds, namely corn and dandelion, whose pollen is reported in the diet of a diversity of bee species in spring and summer (Ammerman et al., 1995; Auclair and Jamieson, 1948; Brian, 1951; Danner et al., 2014; Di Pasquale et al., 2016; Goss, 1968; Henderson et al., 1959; Hogan et al., 1955; Requier et al., 2015; Teper, 2006). Understanding the consequences of tryptophan deficiencies on bees is therefore critical to better manage environmental resources available to them, with the aim of increasing the efficiency of conservation plans and pollination practices.

Here, we investigated how tryptophan deficiencies mimicking those found in corn or dandelion pollen (Auclair and Jamieson, 1948; Goss, 1968) influenced food intake, lifespan and aggressiveness in the buff-tailed bumble bee (*Bombus terrestris*), a major pollinator of crops and wild plants. We exposed micro-colonies of bumble bees to sucrose-based solutions varying in their tryptophan and vitamin B3 content. We predicted that tryptophan deficiencies in the diet would reduce food intake and longevity, and increase aggressiveness, while supplementation in vitamin B3 should buffer these effects (Fengkui et al., 2015; Génissel et al., 2002; Pucilowski and Kostowski, 1983; Velthuis, 1992).

## Materials and Methods

### Bees

We purchased four *B. terrestris* colonies (BioBest, Belgium) from which we built 60 micro-colonies. Each of these micro-colony was composed of 10 workers of unknown age randomly collected in the four mother colonies (N = 600 workers). All workers were marked with colored numbered tags on the thorax for individual identification. Micro-colonies were maintained in bipartite plastic boxes (10 × 16 × 16 cm, see (Kraus et al., 2019)) under controlled conditions: temperature 25-27°C; humidity 35–45%; photoperiod 12 L: 12 D. Bees were fed *ad-libitum* with 50% (w/w) sucrose solution supplemented in the 10 essential amino acids (de Groot, 1953) as described below.

### Artificial diets

Micro-colonies were fed one of four artificial diets (15 micro-colonies/diet and 150 workers/diet): a control diet and three treatment diets with varying concentrations of tryptophan and nicotinamide (Table 1). The control diet was prepared based on previously measured consumptions of *Rubus* pollen by bumble bees (see Table S2 in (Di Pasquale et al., 2013)). *Rubus* is a valuable monofloral source of pollen for honey bees rich in protein and whose content in EAAs matches that of pollens collected by *B. terrestris* (Kriesell et al., 2017; Leonhardt and Blüthgen, 2012; Vanderplanck et al., 2014). Assuming that *B. terrestris* workers assimilate approximately 20% of ingested amino acids from ingested proteins (Stabler et al., 2015) and that their daily intake of pollen under captive conditions is of 0.3 g maximum (Moerman et al., 2017; Vanderplanck et al., 2014), the estimated daily requirement of each EAA was estimated to match the content in each essential amino acid of *Rubus spp*. pollen as described in Table S2 of (Di Pasquale et al., 2013). As EEA content was provided in g/100g of pollen, we multiplied the value by 10 to obtain a value in mg/g. Thus, the daily requirement of workers was assessed as follows:

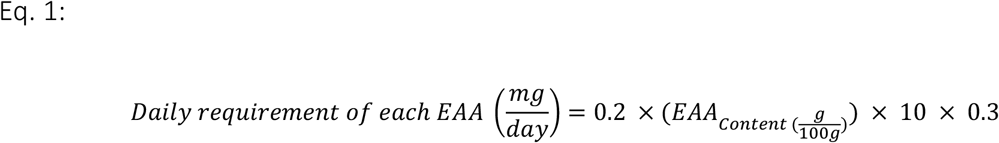

The optimal content, based on *Rubus spp*. pollen, was provided to the control group. The three treatment diets (MT, LT and LTN) had the same content than the control diet for 9 of the EAAs, but a reduced content in tryptophan, with either medium (MT) or low (LT) tryptophan levels. The LTN diet had an identical EAAs composition to the LT diet but was supplemented in nicotinamide (Table 1). Workers of *B. terrestris* ingest on average 1 mL of control diet per day in our laboratory conditions (*preliminary data not shown*) which matches the estimated daily intake in EAAs if bees were fed *Rubus* pollen.

**Table 1.** Concentrations of essential amino acids (in mg/mL) in the diets. Estimated daily consumption of each of the 10 EEAs by bumble bee workers if fed on *Rubus* pollen, based on its composition (Di Pasquale et al., 2013) and daily intake and assimilability of EAAs from pollen (Kriesell et al., 2017; Leonhardt and Blüthgen, 2012; Stabler et al., 2015; Vanderplanck et al., 2014). Control = control diet, mimicking the composition of *Rubus* pollen, MT = medium tryptophan, LT = low tryptophan, and LTN = low tryptophan with nicotinamide supplementation.

| Diet |  | *Rubus* sp. | Control | MT | LT | LTN |
| --- | --- | --- | --- | --- | --- | --- |
| EAAs | Unit | mg/day | mg/mL | mg/mL | mg/mL | mg/mL |
| Valine |  | 0.684 | 0.684 | 0.684 | 0.684 | 0.684 |
| Methionine |  | 0.324 | 0.324 | 0.324 | 0.324 | 0.324 |
| Isoleucine |  | 0.546 | 0.546 | 0.546 | 0.546 | 0.546 |
| Leucine |  | 0.888 | 0.888 | 0.888 | 0.888 | 0.888 |
| Threonine |  | 0.354 | 0.354 | 0.354 | 0.354 | 0.354 |
| Phenylalanine |  | 0.594 | 0.594 | 0.594 | 0.594 | 0.594 |
| Lysine |  | 0.9 | 0.9 | 0.9 | 0.9 | 0.9 |
| Histidine |  | 0.276 | 0.276 | 0.276 | 0.276 | 0.276 |
| Arginine |  | 0.618 | 0.618 | 0.618 | 0.618 | 0.618 |
| Tryptophan |  | 0.162 | 0.162 | 0.005 | 0.0001 | 0.0001 |
| Nicotinamide |  | / | / | / | / | 0.006 |

Diets were prepared in a final volume of 500 mL. They consisted in 50% sucrose solution (w/w) with 0.5% of insect vitamin cocktail (Sigma, Germany), plus the corresponding EAA (Table 1). In order to adjust low EAA concentrations, tryptophan was added to MT, LT and LTN diets from an intermediate solution at 0.05 mg/mL of tryptophan in distilled water. Nicotinamide in LTN was added from a solution of 3mg/mL of nicotinamide in distilled water. Diets were provided in a gravity feeder consisting of a 15 mL tube with two holes at its basis, allowing bumble bees to insert their proboscis and ingest the solution.

### Food intake and survival

We let the bumble bees acclimate to the micro-colony setup during six days. We then monitored the food intake of the workers by renewing all diets every two days with a new cup to collect potential leakage (Kraus et al., 2019). Each vial and cup were weighed before and after being provided to the bees from day 7 (first behavioral trial) to day 18 (end of the experiment), using a precision balance of 1mg (ME103T, Mettler Toledo, Switzerland). Evaporation of the sucrose solution was estimated by placing six gravity feeders in the room and weighing them every two days. Dead bees were recorded and removed daily in each micro-colony. The evaporation and number of living bumble bees were then used to assess individual daily food intake (intake of the sucrose solution in g.bee.day^-1^) and cumulated food intake throughout the experiment. Survival was assessed on 400 bumble bees from 40 micro-colonies (n=10/diet); workers from these micro-colonies did not experience behavioral tests.

Calculation of daily and cumulative food intake per bee was conducted following (Kraus et al., 2019), as shown below. We consider food collected as food ingested, given that there is no trophallaxis in bumble bees and that there was no honey pot through which the sucrose solution could be stored in our microcolonies (Dornhaus and Chittka, 2005; Liebig et al., 1997).

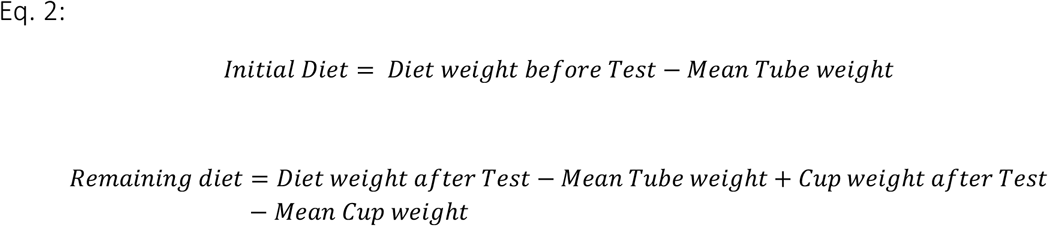

Measure of leakage:

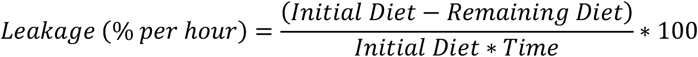

Amount of food ingested:

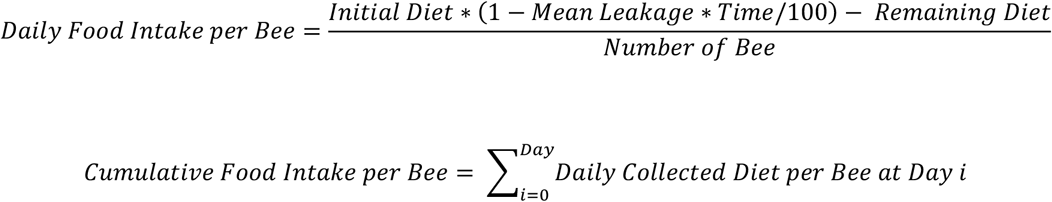

### Behavioral interactions and aggressive behavior

Behavioral assays were conducted on individuals from 20 of the 60 micro-colonies (5 micro-colonies/diet; Table 1) on days 7 and 15. We quantified aggressiveness in dyads of individuals (10 different dyads; Figure 1). Each trial consisted of a 5min acclimation period during which two bumble bees were placed in a petri-dish (10cm diameter) divided in two chambers with a plastic separator. After this period, the separator was removed to allow the bumble bees to interact. Each trial was videotaped during the 5min (Sony Handycam Flash HDR-CX240EB), which started at the removal of the plastic separator. We tested all 10 diet combinations and obtained 2-3 replicates for each combination (see Table S1). We recorded and differentiated total physical and aggressive interactions between workers. Physical interactions included all occurrences with body contact (antennation, head or abdomen contact) between the two workers. Aggressive interactions included biting and body contacts with opened mandibula, which are considered as extreme expressions of aggression, i.e attacks (Amsalem and Hefetz, 2011; Duchateau, 1989; Padilla et al., 2016). After each trial, workers were placed back in their respective micro-colonies.

**Figure 1:**
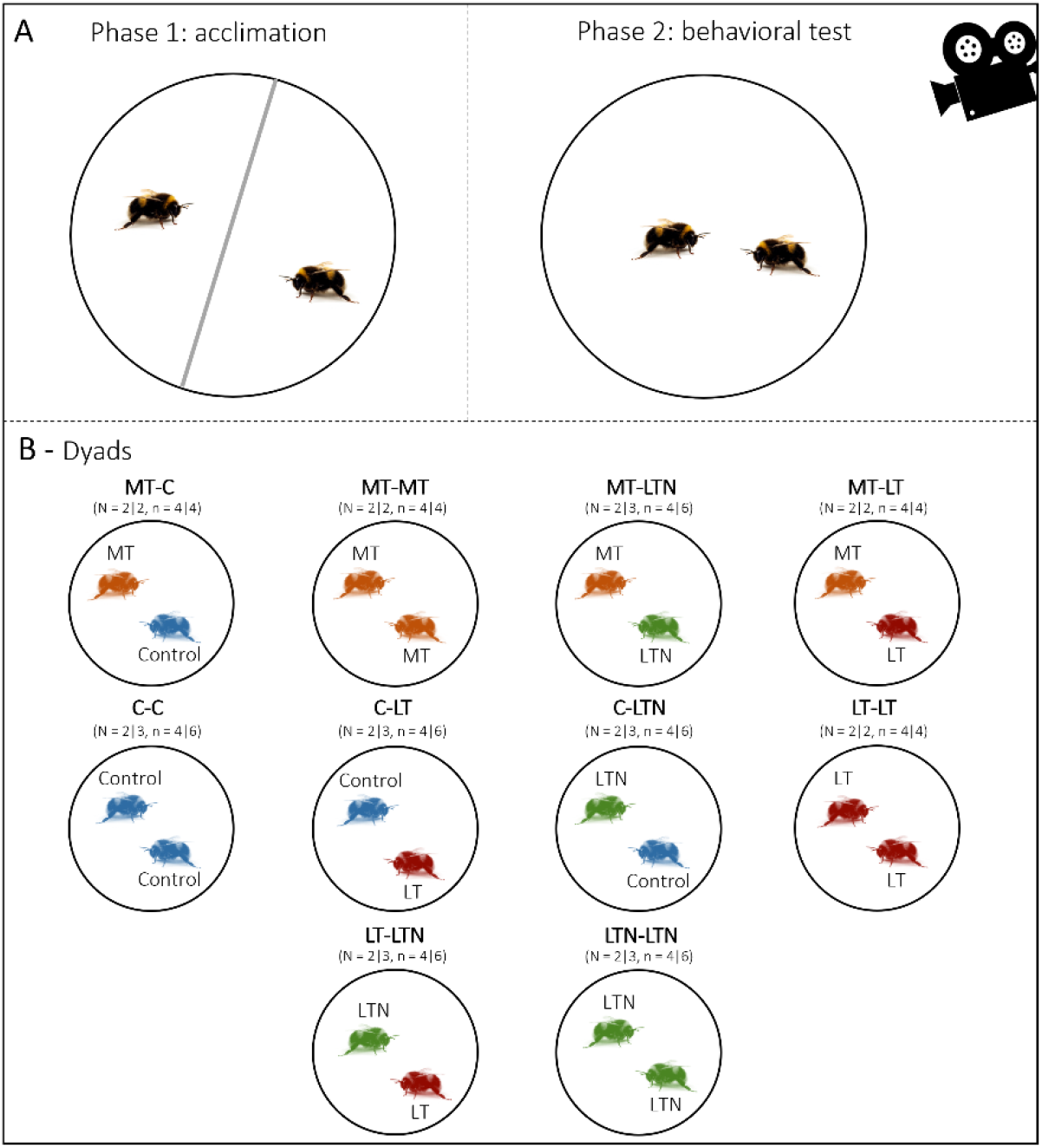
Experimental design used to assess workers behavior. In (A) the two phases of the behavioral test are shown. Phase 1 corresponds to the 5min acclimation period with a plastic separator preventing workers to interact. Phase 2 corresponds to the 5min viteotapped behavioral trial, during which bees were allowed to interact in the petri dish. In (B) the 10 dyads tested are represented. Numbers of dyads (N) and number of bees (n) tested per diet are shown for each trial, respectively (first | second). A total of 92 bees were tested (40 in trial 1, and 52 in trial 2, 21-25 bees/diet).

### Statistical analyses

#### Food intake

We looked at the diet effect on the cumulated food intake from day 7 to day 18 using a linear mixed model (LMM). We included the diet, the day and the diet*day interaction as fixed effects while the micro-colony ID was included as a random effect to control for repeated measures on the same micro-colonies. We also investigated the effect of the diet on the daily food intake (g.bee.day^-1^) on the last day of the experiment (daily food intake on day 18) using a linear model (LM) with the diet as fixed effect.

#### Survival

We conducted a Kaplan-Meier survival analysis to look at the effect of the diet on workers survival from day 0 to day 18. Paired comparisons were conducted using the Log Rank (Mantel-Cox) test. This study was conducted using workers from the 40 micro-colonies that were not used for behavioral tests. An important mortality occurred in all diet groups in the first 24 hours of the experiment, likely because of handling stress following tagging. Workers that died during this timeframe were replaced with new individuals (see Methods).

#### Interactions and aggressive behavior

We used LMMs to investigate diet effects on the number of interactions and number of attacks between workers during the behavioral trials. We included the diet, the date of trial and the interaction between these two variables as fixed effects. The micro-colony ID and colony of origin were added as random effects to control for repeated measures on the same micro-colonies and for potential genetic effects on aggressiveness (i.e., colony of origin). We initially estimated that a minimum of 10 repetitions/dyad or combinations would be necessary to assess a dyad effect. However, due to the high mortality of bees during the first 48 hours of the study and considering that we did not have enough bees to replace all the workers in the micro-colonies destined to behavioral tests, we ended with 2-3 repetitions per dyad (Figure 1). We thus could not test for a dyad*diet interaction effect (e.g., whether control bees reacted differently when confronted to LT, MT or LTN bees).

In all linear models, normality of the residuals was tested using a Kolmogorov-Smirnov test and homogeneity of variances using a Bartlett test. Analyses were conducted using SPSS software (IBM SPSS Statistics for WINDOWS v. 24.0. Armonk, NY: IBM Corp.). Data presented are means ± SEM except when specified otherwise.

## Results

### 1. Tryptophan deficiencies increased food intake

We found no significant effect of the diet on the average individual cumulated food intake between day 7 and day 18 (LMM, F_3;51.5_ = 0.44, p = 0.72, Figure 2A). However, there was an effect of the diet on the average individual daily food intake on day 18 (LM, F_3;31_ = 3.18, p = 0.038, Figure 2B). On the last day of the experiment, bumble bees from the LT and MT diet groups ingested significantly more food than bees from the control diet group (Figure 2B, mean difference of 0.018 ± 0.006 and 0.017 ± 0.007, respectively, t = 2.72, p = 0.011 and t = 2.57, p = 0.015). Bumble bees from the LTN diet group had an intermediate daily food intake (average daily food intake on day 18: C = 0.017 ± 0.005, LT = 0.035 ± 0.004, MT = 0.034 ± 0.005, LTN = 0.026 ± 0.005 g.bee.day^-1^). Tryptophan deficiencies thus increased food intake on the long term and this effect was partially counteracted by nicotinamide supplementation.

**Figure 2:**
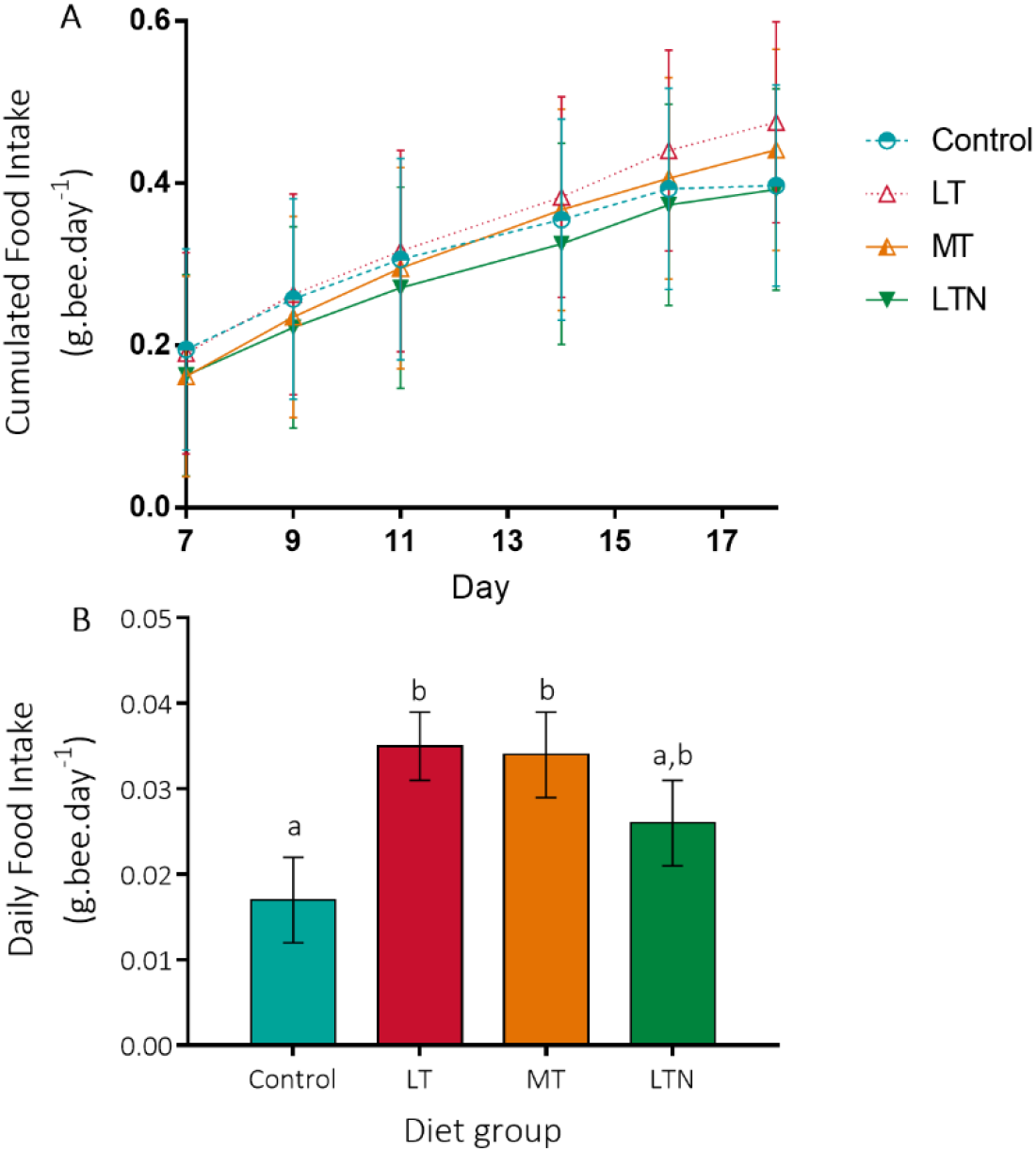
Food intake. (A) Individual cumulated food intake (in g.bee.day-1) from day 7 to day 18, and (B) individual daily food intake on day 18. C = Control, LT = low-tryptophan, MT = medium-tryptophan and LTN = low-tryptophan with nicotinamide supplementation. Different letters represent significant differences between the groups (LM, p<0.05). Means ± SEM are represented.

### 2. Tryptophan deficiencies decreased survival

We found a diet effect on survival (Figure 3). Paired comparisons (Log-rank/Mantel-Cox test) showed that workers fed LT diet had reduced survival compared to workers fed control diet (*X*^2^ = 8.11, df = 3, p = 0.004,). Workers fed MT and LTN diets displayed intermediate survival rates, non-significantly different from that fed control diet (*X*^2^ = 0.90, df = 3, p = 0.34, and *X*^2^ = 2.61, df = 3, p = 0.11). The survival rate of workers fed MT diet, however, differed significantly from that of workers fed LT diet (*X*^2^ = 5.80, df = 3, p = 0.016), while this difference was non-significant between workers fed LTN and LT diets (*X*^2^ = 0.13, df = 3, p = 0.72). This analysis was made without including the data of first 48h since many workers died and were replaced with new ones (see Methods). Including the data from the first 48h provided similar results with an intermediate survival rate for the LTN group (Figure S1 and supplementary results). Tryptophan deficiencies thus reduced survival and this effect was partially counteracted by nicotinamide supplementation.

**Figure 3:**
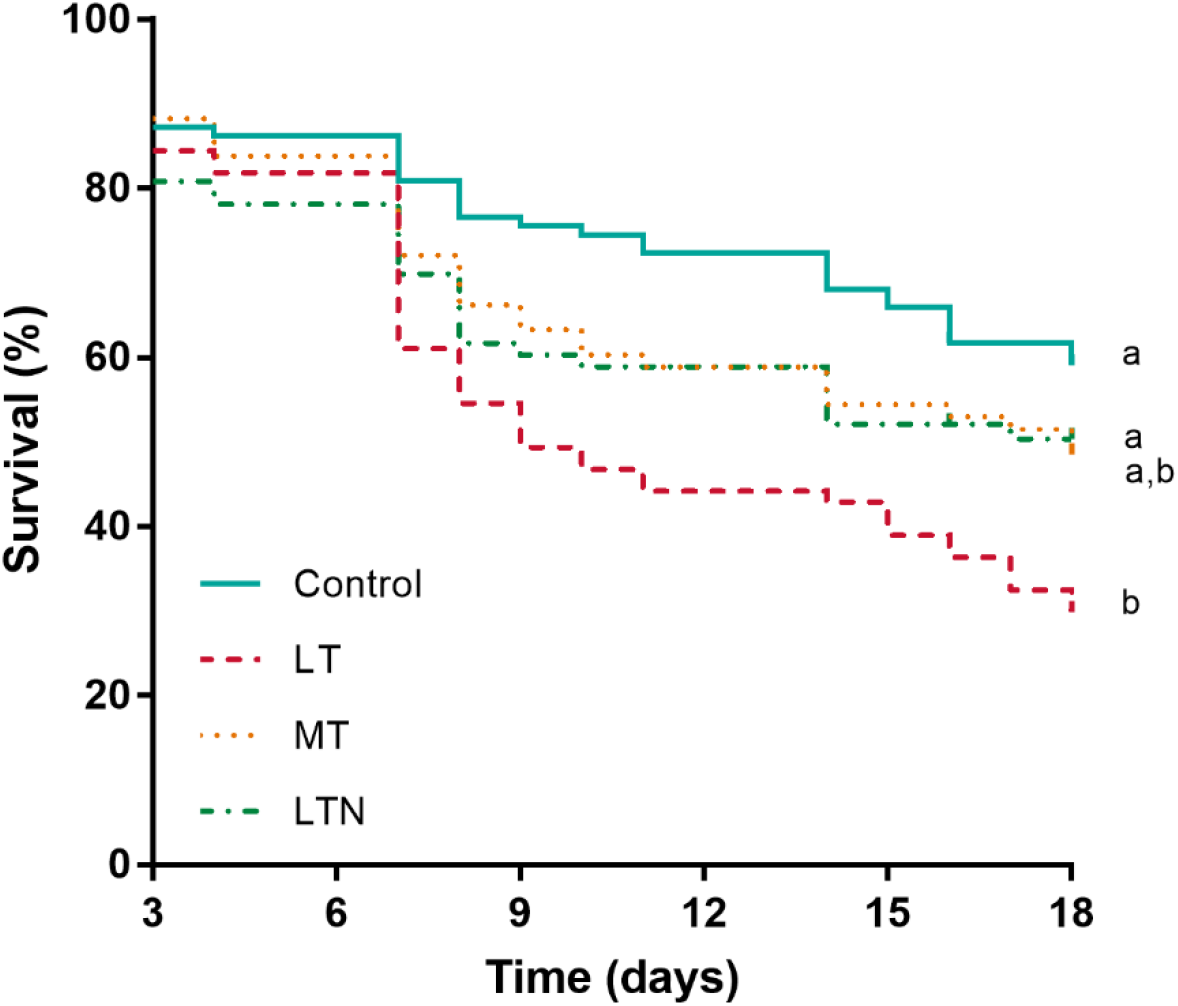
Surviva. Survival rates are shown between day 3 and day 18, considering the high mortality that occurred during the first 48h of the study and the replacement of workers (Survival rates for the entire period are shown on Figure S1). Different letters indicate significant differences between the diet groups (Kaplan-Meier survival curve, p<0.05). LT = low-tryptophan, MT = medium-tryptophan and LTN = low-tryptophan with nicotinamide supplementation.

### 3. Tryptophan deficiencies increased aggressiveness

We found a significant diet*period interaction between (F_3;82_ = 16.329, p < 0.001) on the mean number of physical interactions between workers during the behavioral trials (Figure 4A). On day 7, bumble bees fed LT diet had significantly more interactions with their conspecifics than bumble bees fed the three other diets (p < 0.004, Figure 4A). In contrast, on day 15-16, the mean number of physical interactions was significantly greater in bumble bees fed LTN diet than in bumble bees fed the other diets (p = 0.001, Figure 4A). This could be related to the expression of defensive behaviors by some of the LTN fed bees when exposed to MT and LT fed bees, which were observed in those trials. These defensive behaviors included “prostrate” (one bee is lying on the side, while another bee is involved in an aggressive interaction or an attack) and “escape” (one bee is moving fast in an opposite direction/to create distance when another bee is following or attacking). However, their appearance was too scarce for us to conduct statistical analyses.

**Figure 4:**
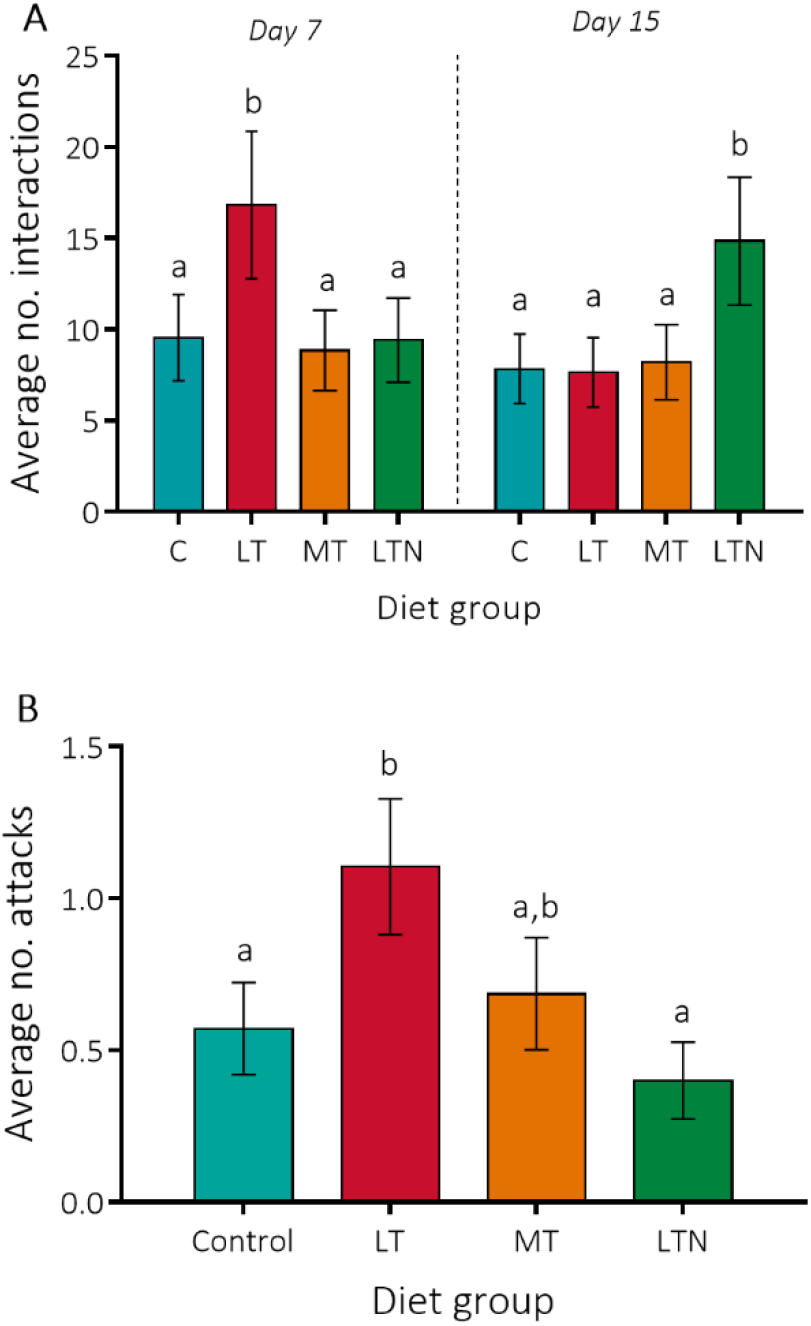
Behavior. (A) Average number of physical interactions recorded between workers at day 7 (n=40) and day 15 of the experiment (n=50, among which 10 dyads were tested on day 16). (B) Average number of attacks recorded from workers fed a given diet. LT = low-tryptophan, MT = medium-tryptophan and LTN = low-tryptophan with nicotinamide supplementation. Different letters highlight a significant difference (LMM, p < 0.05) between the diet groups. Means ± SEM are represented.

We also found a significant effect of diet on the number of attacks (F_3;82_ = 3.017, p = 0.035), and an absence of effects of the period (F_1;82_ = 2.070, p = 0.154) and the interaction between these two variables (F_3;82_ = 0.788, p = 0.504). Bumble bees fed LT diet expressed significantly more attacks than bumble bees fed control and LTN diets (Figure 4B, p=0.035 and p=0.009, respectively). The average number of attacks expressed by bumblebees fed MT diet was intermediate but did not differ significantly from those expressed by bumble bees fed LTN, LT and control diets (Figure 4B, p=0.12, p=0.25 and p=0.57). Tryptophan deficiencies thus increased aggressiveness among workers and this effect was counteracted by nicotinamide supplementation.

## Discussion

Previous studies showed a lack of several essential amino acids can have severe consequences on the reproductive and survival performances of bees (Leonhardt and Blüthgen, 2012; Moerman et al., 2017; Moerman et al., 2016; Ryder et al., 2021; Stabler et al., 2015; Vanderplanck et al., 2014). However, the effects of tryptophan have been overlooked. Here we found a lack of tryptophan is detrimental, as reported in many vertebrates (Baker, 2008; Kantak et al., 1980; Tissier et al., 2017; Walz et al., 2013). Tryptophan-deficient diets increased food intake, aggressiveness and mortality in bumble bees. Importantly, supplementation of vitamin B3, a metabolic product of tryptophan that must be obtained from the diet, buffered these effects.

### Dietary tryptophan deficiencies induced an increase in food intake, partially buffered by vitamin B3 supplementation

Bumble bee workers fed LT and MT diets tended to ingest more food overall than workers fed control diet throughout the experiment, however, this difference was non-significant (cumulated food intake, Figure 1A). Nonetheless, this resulted in significant differences in the daily food intake at the end of the experiment between the diet groups. On day 18, workers from the LT and MT diet groups ingested twice as much food than bees from the control group, with workers from the LTN group displaying intermediate food intakes. These effects are opposite to what we initially predicted based on a previous study in honey bees, where a sub-optimal content in tryptophan in diet led to reduced food intake (Fengkui et al., 2015). In that study, however, the authors sought to identify the optimal intake of tryptophan by the honey bee, using diets containing between 9-14 mg/g of tryptophan, and composed of a mix between rape pollen and sucrose (Fengkui et al., 2015). This is much higher than the tryptophan content of our LT and MT diets, where we sought to mimic the deficiencies found in corn and dandelion pollen, with tryptophan content ≤1mg/g (Auclair and Jamieson, 1948; Goss, 1968; Hsu et al., 2021). In our study, bumble bees may have tended to increase their daily food intake to compensate for the deficiency in tryptophan, trying to meet their daily requirement in this EAA, as observed in ants (Csata et al., 2020). Alternatively, considering that tryptophan (as a precursor of serotonin) is known to control appetite (French et al., 2014), the strong deficiency of tryptophan in the LT and MT diets could have led to a loss in the down regulation of appetite and a consequent increase in food intake (French et al., 2014). Since bumble bees rely less on nectar than honey bees, and collect pollen with significantly higher content in amino acids (Leonhardt and Blüthgen, 2012), their daily requirements (optimum) and responses when below or above this optimum may vary, as well as the downregulation from serotonin on food intake.

### Tryptophan deficiencies reduced workers survival, which was not fully counteracted by vitamin B3 supplementation

Tryptophan deficiencies reduced worker survival compared to the control group, as previously observed in honey bees fed with a tryptophan-deficient diet (Fengkui et al., 2015) or fed with maize (Hocherl et al., 2012; Velthuis, 1992) known to be deficient in tryptophan (Goss, 1968). These negative effects of tryptophan deficiencies thus could not be prevented by increased food intakes. However, vitamin B3 supplementation tended to increase bumble bee survival compared to the LT diet group from day 10 forward although this difference was non-significant. This suggests the vitamin B3 supplementation of 0.006 mg/mL in sucrose solution was too low in our study and that higher intakes of vitamin B3 are necessary to promote survival levels like that of bumble bees fed control diets, especially in a tryptophan-deficient diet. Given that the optimal tryptophan and vitamin B3 intakes for bees other than the honey bees (Fengkui et al., 2015) are unknown, future studies are needed to quantify precise daily intakes and correlate them with performance traits (e.g., survival and reproduction).

### Vitamin B3 supplementation counteracted the negative effects of tryptophan deficiency on workers aggressiveness

Our results show tryptophan deficiencies also led to increased aggressiveness in worker bumble bees, with more attacks in workers fed LT diet compared to workers fed control and LTN diets, while workers fed MT diet displayed intermediate values. Although vitamin B3 supplementation was presumably too low to significantly counteract the detrimental effects of tryptophan deficiencies on survival, it had a marked effect on behavior. A supplementation in vitamin B3 of 0.006 mg/mL of sucrose solution significantly increased the number of interactions between bees on day 15-16 compared to the other diets, and significantly reduced the occurrences of aggressive interactions (attacks) between workers compared to those of the LT diet. This could be explained by the expression of defensive behaviors from bumble bees fed LTN, when confronted to bumble bees fed LT or MT diets, as occasionally observed during the second trials. Bees fed LTN diet also displayed the lowest number of aggressive interactions when confronted to workers from other micro-colonies, which was twice as low as observed in the LT diet. These behavioral modifications might either be a consequence of modifications in the serotonin pathway or resulting from benefits of vitamin B3 on neuron health and functioning. Indeed, serotonin is known to modulate aggressiveness in many species (Bubak et al., 2020; Pucilowski and Kostowski, 1983) and defensive behaviors in the honey bee (Hunt, 2007). Its synthesis could be related to diet content in vitamin B3 through tryptophan, which is a precursor of both of these molecules (de Arruda et al., 2013; Kohlmeier, 2015). Alternatively, vitamin B3 is known to be essential for cell functioning and ATP synthesis: it can be decomposed into two molecules, nicotinic acid and nicotinamide, which is at the base of the synthesis of the coenzyme NAD (Wan, Moat, & Anstey, 2011). In mammals, NAD serves as neural modulator and may regulate many behaviors, including anti-predatory behaviors (Richards et al., 1983). The pathways by which tryptophan deficiency and vitamin B3 supplementation modulated the behavior of bumble bees remain to be further investigated, but the role of these nutrients in modulating aggressiveness and social interactions thus appear essential. We therefore need ecologically driven studies on bee nutrition that will assess pollen content in tryptophan and vitamin B3 and how daily intakes of these nutrients can impact bee performance, as has been done for the 9 other EAAs and other essential nutrients (Arien et al., 2015; Conroy et al., 2016; Kriesell et al., 2017; Moerman et al., 2017; Moerman et al., 2016; Ryder et al., 2021; Vanderplanck et al., 2014; Vaudo et al., 2016).

### Ecological and fitness-related perspectives

Pollen of some widely-distributed plants, such as corn and dandelion, are known to be deficient in tryptophan and some other essential amino acids for bees (Auclair and Jamieson, 1948; Goss, 1968). Honey bees fed maize or dandelion pollen showed strongly reduced survival rates as well as reduced rearing capacity and reproductive success compared with bees fed other pollen (Frias et al., 2015; Hocherl et al., 2012; Roulston and Cane, 2000; Standifer et al., 1980; Velthuis, 1992). Consumption of maize pollen also impaired the development of the hypopharyngeal gland acini and reduced vitellogenin gene expression in nurses (Di Pasquale et al., 2016). Furthermore, bumble bees fed dandelion pollen displayed very high rates of oophagy and larval ejection (100%), likely because of a deficiency in an essential amino acid (Génissel et al., 2002). Although the direct link between tryptophan deficiencies and reduced performances has not been established in these studies, together with our findings, these observations echo what has been reported in a vertebrate, where corn consumption and associated tryptophan and vitamin B3 deficiencies led to abnormal behavior, leading to 95% of maternal infanticides as well as cannibalism, counteracted by vitamin B3 supplementation (Tissier et al., 2017). Given the preponderance of dandelion and corn in terrestrial landscapes, this could have important ecological consequences for a wide diversity of bee species, known to collect these pollens (Brian, 1951; Requier et al., 2015; Teper, 2006), by affecting not only their survival and aggressive behavior but also their reproduction (Génissel et al., 2002; Vanderplanck et al., 2020). Ultimately, the more detailed understanding of the nutritional requirements of bees at the level of micronutrients will help better managing environmental resources made available to pollinators, for conservation and pollination purposes.

## Acknowledgements

We thank Audrey Baylet for her help in collecting part of the data. Many thanks to Cristian Pasquaretta and Maxime Choblet for their help with collecting preliminary data on related experiment allowing to publish this paper.

## Author Contributions

M.L.T. conceived the theoretical framework and design of this study, with feedback from S.K. and M.L. Data collection was supervised by S.K. and T.G-M. M.L.T. conducted statistical analyses and wrote the first draft of the manuscript with edits from all other authors.

## Data and materials availability

All data will be made available on OSF.

## References

Ammerman, C. B., Baker, D. P. and Lewis, A. J. (1995). Bioavailability of nutrients for animals: Amino acids, minerals, vitamins: Elsevier.

Amsalem, E. and Hefetz, A. (2011). The effect of group size on the interplay between dominance and reproduction in Bombus terrestris. PLoS One 6, e18238.

Arien, Y., Dag, A., Zarchin, S., Masci, T. and Shafir, S. (2015). Omega-3 deficiency impairs honey bee learning. Proceedings of the National Academy of Sciences 112, 15761–15766.

Auclair, J. L. and Jamieson, C. A. (1948). A qualitative analysis of amino acids in pollen collected by bees. Science 108, 357–358.

Baker, D. H. (2008). Animal models in nutrition research. The Journal of nutrition 138, 391–396.

Brian, A. D. (1951). The pollen collected by bumble-bees. The Journal of Animal Ecology, 191–194.

Brodschneider, R. and Crailsheim, K. (2010). Nutrition and health in honey bees. Apidologie 41, 278–294.

Bubak, A. N., Watt, M. J., Yaeger, J. D. W., Renner, K. J. and Swallow, J. G. (2020). The stalk-eyed fly as a model for aggression - is there a conserved role for 5-HT between vertebrates and invertebrates? Journal of Experimental Biology 223, jeb132159.

Conroy, T. J., Palmer-Young, E. C., Irwin, R. E. and Adler, L. S. (2016). Food limitation affects parasite load and survival of Bombus impatiens (Hymenoptera: Apidae) infected with Crithidia (Trypanosomatida: Trypanosomatidae). Environmental Entomology 45, 1212–1219.

Csata, E., Gautrais, J., Bach, A., Blanchet, J., Ferrante, J., Fournier, F., Lévesque, T., Simpson, S. J. and Dussutour, A. (2020). Ant Foragers Compensate for the Nutritional Deficiencies in the Colony. Current Biology 30, 135–142.e4.

Danner, N., Härtel, S. and Steffan-Dewenter, I. (2014). Maize pollen foraging by honey bees in relation to crop area and landscape context. Basic and Applied Ecology 15, 677–684.

de Arruda, V. A. S., Pereira, A. A. S., de Freitas, A. S., Barth, O. M. and de Almeida-Muradian, L. B. (2013). Dried bee pollen: B complex vitamins, physicochemical and botanical composition. Journal of Food Composition and Analysis 29, 100–105.

de Groot, A. P. (1953). Protein and amino acid requirements of the honeybee (Apis mellifica L.).

Di Pasquale, G., Alaux, C., Le Conte, Y., Odoux, J. F., Pioz, M., Vaissiere, B. E., Belzunces, L. P. and Decourtye, A. (2016). Variations in the Availability of Pollen Resources Affect Honey Bee Health. PLoS One 11, e0162818.

Di Pasquale, G., Salignon, M., Le Conte, Y., Belzunces, L. P., Decourtye, A., Kretzschmar, A., Suchail, S., Brunet, J.-L. and Alaux, C. (2013). Influence of Pollen Nutrition on Honey Bee Health: Do Pollen Quality and Diversity Matter? PLoS One 8, e72016.

Dornhaus, A. and Chittka, L. (2005). Bumble bees (Bombus terrestris) store both food and information in honeypots. Behavioral Ecology 16, 661–666.

Duchateau, M. J. (1989). Agonistic Behaviours in Colonies of the Bumblebee Bombus terrestris. Journal of Ethology 7, 141–151.

Fengkui, Z., Baohua, X., Ge, Z. and Hongfang, W. (2015). The Appropriate Supplementary Level of Tryptophan in the Diet of Apis mellifera (Hymenoptera: Apidae) Worker Bees. Journal of Insect Science 15.

French, A. S., Simcock, K. L., Rolke, D., Gartside, S. E., Blenau, W. and Wright, G. A. (2014). The role of serotonin in feeding and gut contractions in the honeybee. Journal of Insect Physiology 61, 8–15.

Frias, B. E. D., Barbosa, C. D. and Lourenço, A. P. (2015). Pollen nutrition in honey bees (Apis mellifera): impact on adult health. Apidologie 47, 15–25.

Génissel, A., Aupinel, P., Bressac, C., Tasei, J.-N. and Chevrier, C. (2002). Influence of pollen origin on performance of Bombus terrestris micro-colonies. Entomologia Experimentalis et Applicata 104, 329–336.

Goss, J. A. (1968). Development, physiology, and biochemistry of corn and wheat pollen. The Botanical Review 34, 333–359.

Goulson, D., Nicholls, E., Botías, C. and Rotheray, E. L. (2015). Bee declines driven by combined stress from parasites, pesticides, and lack of flowers. Science 347.

Hegyi, J., Schwartz, R. A. and Hegyi, V. (2004). Pellagra: dermatitis, dementia, and diarrhea. International journal of dermatology 43, 1–5.

Henderson, L., Someroski, J. F., Rao, D., Wu, P.-H. L., Griffith, T. and Byerrum, R. U. (1959). Lack of a tryptophan-niacin relationship in corn and tobacco. Journal of Biological Chemistry 234, 93–95.

Hocherl, N., Siede, R., Illies, I., Gatschenberger, H. and Tautz, J. (2012). Evaluation of the nutritive value of maize for honey bees. J Insect Physiol 58, 278–85.

Hogan, A., Gillespie, G., Koçtürk, O., O’dell, B. and Flynn, L. M. (1955). The percentage of protein in corn and its nutritional properties. The Journal of nutrition 57, 225–239.

Hsu, P. S., Wu, T. H., Huang, M. Y., Wang, D. Y. and Wu, M. C. (2021). Nutritive Value of 11 Bee Pollen Samples from Major Floral Sources in Taiwan. Foods 10, 2229.

Hunt, G. J. (2007). Flight and fight: A comparative view of the neurophysiology and genetics of honey bee defensive behavior. Journal of Insect Physiology 53, 399–410.

Kantak, K. M., Hegstrand, L. R. and Eichelman, B. (1980). Dietary tryptophan modulation and aggressive behavior in mice. Pharmacology Biochemistry and Behavior 12, 675–679.

Klein, S., Cabirol, A., Devaud, J.-M., Barron, A. B. and Lihoreau, M. (2017). Why bees are so vulnerable to environmental stressors. Trends in ecology & evolution 32, 268–278.

Kohlmeier, M. (2015). Nutrient metabolism: structures, functions, and genes: Academic Press.

Kraus, S., Gómez-Moracho, T., Pasquaretta, C., Latil, G., Dussutour, A. and Lihoreau, M. (2019). Bumblebees adjust protein and lipid collection rules to the presence of brood. Current Zoology 65, 437–446.

Krehl, W., Teply, L., Sarma, P. and Elvehjem, C. (1945). Growth-retarding effect of corn in nicotinic acid-low rations and its counteraction by tryptophane. Science 101, 489–490.

Kriesell, L., Hilpert, A. and Leonhardt, S. D. (2017). Different but the same: bumblebee species collect pollen of different plant sources but similar amino acid profiles. Apidologie 48, 102–116.

Leonhardt, S. D. and Blüthgen, N. (2012). The same, but different: pollen foraging in honeybee and bumblebee colonies. Apidologie 43, 449–464.

Liebig, J., Heinze, J. and Hölldobler, B. (1997). Trophallaxis and Aggression in the Ponerine Ant, Ponera coarctata: Implications for the Evolution of Liquid Food Exchange in the Hymenoptera. Ethology 103, 707–722.

Meisinger, D. J. and Speer, V. (1979). Tryptophan requirement for reproduction in swine. Journal of animal science 48, 559–569.

Moerman, R., Vanderplanck, M., Fournier, D., Jacquemart, A. L. and Michez, D. (2017). Pollen nutrients better explain bumblebee colony development than pollen diversity. Insect Conservation and Diversity 10, 171–179.

Moerman, R., Vanderplanck, M., Roger, N., Declèves, S., Wathelet, B., Rasmont, P., Fournier, D. and Michez, D. (2016). Growth rate of bumblebee larvae is related to pollen amino acids. Journal of Economic Entomology 109, 25–30.

Padilla, M., Amsalem, E., Altman, N., Hefetz, A. and Grozinger, C. M. (2016). Chemical communication is not sufficient to explain reproductive inhibition in the bumblebee Bombus impatiens. Royal Society Open Science 3, 160576.

Pucilowski, O. and Kostowski, W. (1983). Aggressive behaviour and the central serotonergic systems. Behavioural Brain Research 9, 33–48.

Requier, F., Odoux, J.-F., Tamic, T., Moreau, N., Henry, M., Decourtye, A. and Bretagnolle, V. (2015). Honey bee diet in intensive farmland habitats reveals an unexpectedly high flower richness and a major role of weeds. Ecological Applications 25, 881–890.

Richards, C., Snell, C. and Snell, P. (1983). Nicotinamide adenine dinucleotide depresses synaptic transmission in the hippocampus and has specific binding sites on the synaptic membranes. British journal of pharmacology 79, 553–564.

Roulston, T. H. and Cane, J. H. (2000). Pollen nutritional content and digestibility for animals. Plant Systematics and Evolution 222, 187–209.

Ryder, J. T., Cherrill, A., Thompson, H. M. and Walters, K. F. A. (2021). Lower pollen nutritional quality delays nest building and egg laying in Bombus terrestris audax micro-colonies leading to reduced biomass gain. Apidologie 52, 1033–1047.

Stabler, D., Paoli, P. P., Nicolson, S. W. and Wright, G. A. (2015). Nutrient balancing of the adult worker bumblebee (Bombus terrestris) depends on the dietary source of essential amino acids. The Journal of experimental biology 218, 793–802.

Standifer, L., McCaughey, W., Dixon, S., Gilliam, M. and Loper, G. (1980). Biochemistry and microbiology of pollen collected by honey bees (Apis mellifera L.) from almond, Prunus dulcis. II. Protein, amino acids and enzymes. Apidologie 11, 163–171.

Teper, D. (2006). Food plants of Bombus terrestris as determined by pollen analysis of faeces. Journal of Apicultural Science 50, 101–108.

Tissier, M. L., Handrich, Y., Dallongeville, O., Robin, J.-P. and Habold, C. (2017). Diets derived from maize monoculture cause maternal infanticides in the endangered European hamster due to a vitamin B3 deficiency. Proceedings of the Royal Society B: Biological Sciences 284, 20162168.

Vanbergen, A. J. and Initiative, T. I. P. (2013). Threats to an ecosystem service: pressures on pollinators. Frontiers in Ecology and the Environment 11, 251–259.

Vanderplanck, M., Gilles, H., Nonclercq, D., Duez, P. and Gerbaux, P. (2020). Asteraceae Paradox: Chemical and Mechanical Protection of Taraxacum Pollen. Insects 11.

Vanderplanck, M., Moerman, R., Rasmont, P., Lognay, G., Wathelet, B., Wattiez, R. and Michez, D. (2014). How does pollen chemistry impact development and feeding behaviour of polylectic bees? PLoS One 9, e86209.

Vaudo, A. D., Patch, H. M., Mortensen, D. A., Tooker, J. F. and Grozinger, C. M. (2016). Macronutrient ratios in pollen shape bumble bee (Bombus impatiens) foraging strategies and floral preferences. Proceedings of the National Academy of Sciences 113, E4035–42.

Velthuis, H. H. W. (1992). Pollen Digestion and the Evolution of Sociality in Bees. Bee World 73, 77–89.

Walz, J. C., Stertz, L., Fijtman, A., dos Santos, B. T. and Almeida, R. M. M. d. (2013). Tryptophan diet reduces aggressive behavior in male mice. Psychology & Neuroscience 6, 397–401.

Wan, P., Moat, S. and Anstey, A. (2011). Pellagra: a review with emphasis on photosensitivity. British Journal of Dermatology 164, 1188–1200.

